# Neural Decoding of Inferior Colliculus Multiunit Activity for Sound Category identification with temporal correlation and deep learning

**DOI:** 10.1101/2022.08.24.505211

**Authors:** F. Özcan, A. Alkan

## Abstract

Natural sounds are easily perceived and identified by humans and animals. Despite this, the neural transformations that enable sound perception remain largely unknown. Neuroscientists are drawing important conclusions about neural decoding that may eventually aid research into the design of brain-machine interfaces (BCIs). It is thought that the time-frequency correlation characteristics of sounds may be reflected in auditory assembly responses in the midbrain and that this may play an important role in identification of natural sounds. In our study, natural sounds will be predicted from multi-unit activity (MUA) signals collected in the inferior colliculus. The temporal correlation values of the MUA signals are converted into images. We used two different segment sizes and thus generated four subsets for the classification. Using pre-trained convolutional neural networks (CNNs), features of the images were extracted and the type of sound heard was classified. For this, we applied transfer learning from Alexnet, GoogleNet and Squeezenet CNNs. The classifiers support vector machines (SVM), k-nearest neighbour (KNN), Naive Bayes and Ensemble were used. The accuracy, sensitivity, specificity, precision and F1 score were measured as evaluation parameters. Considering the trials one by one in each, we obtained an accuracy of 85.69% with temporal correlation images over 1000 ms windows. Using all trials and removing noise, the accuracy increased to 100%.

## I. INTRODUCTION

Neuroscientists are seeking to better understand how areas of the brain relate to the external world. Sensory neurons are activated by sensory stimuli such as vision, sound, smell and touch from the outside world. This stimulus information is encoded with spikes (action potentials) and transmitted to the nervous system. One of the goals of neural decoding is to be able to control brain-computer interface devices ([1]- Lazarevich et al, 2020). For this purpose, it is necessary to recognise the code of neural data. The relationship between neural activity and the environment can be better described through neural decoding. In neuroscience, researchers try to learn how the neuronal response (neuronal coding) is encoded. In the decoding phase, the stimulus is determined from the neural activity. For example, the stimulus (image, sound, odour concentration,…) will be predicted from the resulting spike sequence, functional magnetic resonance image (fMRI), electrocorticograph (EcoG), magnetoencephalogram (MEG), calcium imaging, local field potential (LFP), spikes of simple-unit activity (SUA), multi-unit activity (MUA). It is essential to compare the results of studies conducted under different experimental conditions to increase the accuracy of decoding.

Extracellular recording is one of the most widely used electrophysiological techniques for neuroscience studies and for medical applications (e.g. brain-machine interface). The most invasive recordings involve placing electrodes in the brain and recording voltages ([2]- Livezey et al., 2020). Multineuronal extracellular recordings are performed with the access of several probes (multi-electrode arrays) to the brain region. Each individual electrode measures the electrical activity of a neuron, as well as that of neighbouring neurons ([3]- Lefebvre et al., 2018). Decoding studies can be from a single neuron or from a particular brain area (neural network).

Four types of signals can be derived from the acquired data (raw data) by different signal processing steps, as shown in Figure 1. The raw neural recordings are composed of two main components: the local field potential (LFP) and the action potentials (spikes). The spikes can be classified into two types of signals, namely single-unit activity (SUA) and multi-unit activity (MUA). MUA refers to all spikes detected (without spike sorting) from a set of neurons. The MUA has a lower spatial range than the LFP but higher than the unit activity. A bandpass filter is used on the raw neuronal signal to extract the spike waveforms ([4]- Ahmadi et al., 2021). When the threshold crossing in the MUA signal, which is the action potential sequence, is removed, an analog multi-unit activity (aMUA) is obtained. This signal is treated as a time series.

**Figure 1 :**
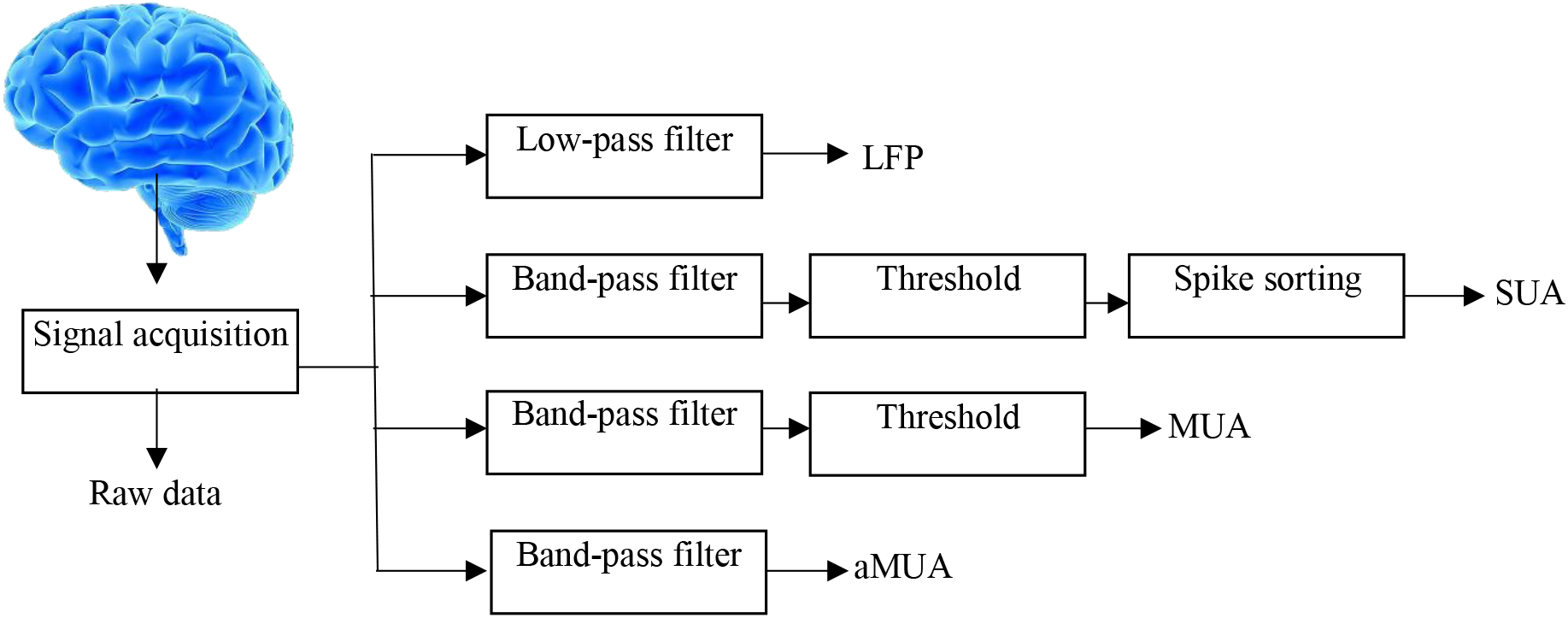
Neuronal signal processing: From raw neural signal acquisition to processing for obtaining different neural signals: Local Field Potential (LFP), Single-Unit Activity (SUA), Multi-Unit Activity (MUA), aMUA analog Multi-Unit Activity (aMUA).

The auditory neural pathway is apparently more complex than the visual one, due to the greater number of synaptic transmissions between the sensory system and the cortex. In addition, signal transmission systems from one nucleus to another in the brain are more frequent in hearing than in the visual system ([5]- Bear et al., 2016). The auditory signal transformations that occur in the brainstem originate from information from spiral ganglion neurons in the cochlea (Figure 2).

**Figure 2:**
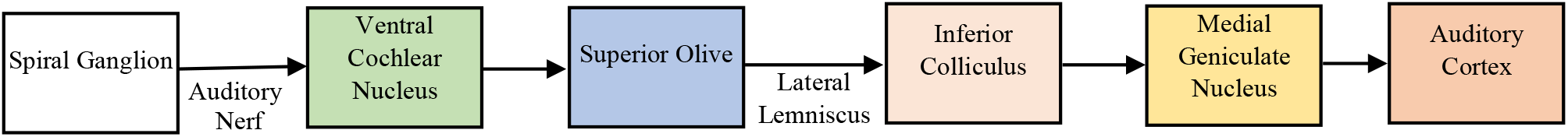
Auditory pathways. Auditory signals converted in the cochlea are transmitted to the auditory cortex via various neural pathways from sensory neurons in the spiral ganglion ([5]- Bear et al., 2016).

Neurons in the auditory system are involved in decoding the duration, intensity and frequency of sound heard ([6]- Richard et al., 2019). Frequency specificity is a property of the neurons in each relay present from the cochlea to the cortex. How is the frequency of the stimulus represented in the nervous system? The active receptor cells in question are not the same depending on the frequency of the sound stimulus. In the brainstem, neurons have properties that allow them to accurately represent different sound frequencies. At very low frequencies, phase correlation is the encoding mechanism; intermediate frequency encoding includes both tonotopy and phase correlation; and at higher frequencies, tonotopy provides a good representation of sound frequencies ([5]- Bear et al., 2016). The systematic organisation of sound frequencies encoded in an auditory structure is called tonotopy. Because of the tonotopy that is found in the whole auditory sensory system, the location of active neurons in the auditory nuclei is an indicator of the frequency of sound and its modulation characteristics ([7]- Downer et al., 2015). Some cells in the cochlear nuclei are particularly sensitive to sounds that vary in frequency over time ([5]- Bear et al., 2016).

The inferior colliculus (IC) is an intermediate level structure located in the main auditory nucleus of the midbrain. The IC receives ascending and converging auditory information from different nuclei in the brainstem. It has the ability to consolidate information which is then relayed to the cortex. Areas of tonotopically organised frequency responses, varying from low to high frequencies with the depth of neuronal recording, are present in the main nucleus of the IC. This is called the tonotopic organisation of the recording area ([8]-Sadeghi et al., 2019). In the inferior colliculus (IC), sounds are broken down into higher order acoustic features such as temporal and spectral modulations. Neurons in the inferior colliculus also respond selectively to the spectro-temporal structure of the sound, rather than being excited when a frequency is simply present in the sound ([7]- Downer et al., 2015).

What is a natural sound, and what types of neural encoding are responsible for recognising natural sounds? The high-level neural processing that enables the recognition of natural sounds is not yet fully understood. It is the structural complexity of these sounds that is responsible. In the case of natural sounds, the spectro-temporal modulations are highly varied and structured, and the envelopes are correlated in frequency and time ([9]- Chen et al., 2012). Cathegories of stationary natural sounds with high-level acoustic structure, such as random sounds from bird chorus, crackling fire, running water, outdoor crowd, and rattling snake sounds were used in this study ([10]-Zhai et al., 2020). Collectively, data we want to use, suggest that stimulus-driven correlations between sets of neurons in the CI can provide an imprint for downstream neurons to recognise and classify natural sounds into categories ([8]-Sadeghi et al., 2019).

On the other hand, for a sound to be recognised by the brain, it must be heard for a certain amount of time. This duration is about 1 to 2 seconds ([8]-Sadeghi et al., 2019). The temporal summation of the messages transmitted by the different neurons will allow to code the intensity and duration of the stimulus ([6]-Richard et al., 2019).

Today, in many machine learning studies, deep learning is the most popular used method. In neuroscience, deep learning has proven to be an important tool for increasing the accuracy and flexibility of neural decoding in a variety of fields ([2]- Livezey et al., 2020). Deep learning techniques can also be considered effective with neural data converted into images.

### 1.1 PURPOSE OF THE WORK

Humans can easily distinguish natural sounds. But this task can be particularly difficult for people with hearing impairments. In order to find a solution to these difficulties, we need to develop the following points:

- Due to the complexity of the physical structure of sounds, the acoustic characteristics and neural coding that allow the recognition of sounds and the formation of corresponding perceptual categories are largely unknown ([8]-Sadeghi et al., 2019).
- When noise is present in neural signals, decoding from the auditory system is difficult.
- Sound categorization performance is highly dependent on the duration of the sound for human listeners (1-2 s). The significant time intervals of a few seconds needed for the perceptual decision process could rather reflect an accumulation process.

In the auditory midbrain, the activity of neural ensembles reveals highly structured correlations. To determine whether neural correlations are affected by the correlation structure of natural sounds and whether this could potentially contribute to sound recognition, Sadeghi et al. use multi-channel neural recordings in the CI ([8]- Sadeghi vd, 2019). To propose an other way to verify this proposal that will allow us to identify, via envelope correlation features, the sounds heard and to improve accuracy results, we have to apply our most advanced and faster computer algorithms to neural decoding.

To see noise effect on the identification of sounds, we will process neural data with and without noise.

From the perspective of neural coding, temporal resolution refers to the period of time over which correlations are computed by neurons. In contrast, the accumulation time represents the period of time during which a neural population reads the correlation values to identify sounds ([8]-Sadeghi et al., 2019). Is it possible that the auditory system integrate sounds on particularly short time. Rather than working only on long time scales, time windows of 250 ms is also used alongside 1000ms windows (corresponding to a 1/2 octave scale) to identify sounds.

Since the duration of the sound heard is very important for identification, it is necessary to highlight the temporal characteristics of neural coding. What extent temporal correlations individually contribute to sound recognition ?

Decoding of neural data using auditory stimulation was only seen in limited numbers in our literature review, and the success rates of these studies were generally relatively low. The implementation of machine learning, deep learning and image processing techniques, which are frequently in use today, will allow for successful results. Our study will be based on the estimation of the auditory stimulus from the neural data, and thus on neural signal decoding. In order to demonstrate the importance of temporal correlation of neural signals, we will apply deep learning methods.

In our study, aMUA signals will be converted into temporal correlation images. With deep learning methods, we will classify neuronal data according to the natural sounds in stimulus, taking into account the temporal correlation, the noise and the temporal resolution. In order to understand neural coding, the objective is to obtain convincing results in the prediction of stimuli with fast techniques using different deep learning models.

Ultimately, this work will help neuroscientists, doctors and engineers to develop solutions for the hearing impaired. In our study, our goal will be to apply deep learning to a field of neuroscience with a multidisciplinary approach. Will deep learning, which comes from the original neural network model, allow us this time to better understand neural data in this context?

### 1.2 RELATED WORK

Machine learning and deep learning techniques have been applied to predict visual stimuli from neural data. However, decoding neural data obtained from an auditory or olfactory stimulus has not been fully studied in the literature, and the success rates of existing studies have remained low. In this study, neural data where the stimulus is sound will be used. Studies similar to the one we have planned are summarised below. In general, different types of data were used and results were obtained with different methods.

Saari et al. conducted a study in 2018, where 18 musicians and 18 non-musicians were played 3 different musical sounds and 116 fMRI images were obtained. A success rate of 0.77 was obtained with linear regression or linear discriminant analysis ([11]-Saari et al., 2018). Bianco et al, in there study intitled “Evaluation of Auditory Cortex Tonotopical Organization in Rats Through Principal Component Analysis”, studied the patterns of neurons in response to ultrasonic vocalisations, in order to find the area of activation corresponding to the perception of vocalisations in rats. By exploiting local field potential (LFP) signal classifications, methods based on statistical analysis were used for this study ([12]-Bianco et al., 2017). Moses et al, in their study entitled “Real-time decoding of question-and-answer speech dialogue using human cortical activity”, classified the EcoG signals collected by 128 electrodes in a question-answer experiment applied to 3 people. A success rate of 76% was obtained for 9 questions and 61% for the answer given by 25 subjects ([13]-Moses et al., 2019).

In addition, a few case studies on decoding neural data with artificial intelligence methods have been conducted. Livezey and Glaser (2020), in their study entitled “Deep Learning approaches for neural decoding: from CNNs to LSTMs and spikes to fMRI”, examined deep learning approaches for neural decoding. To increase success and improve processing capacity in neural decoding, deep learning has proven to be an effective tool in a wide variety of tasks. In the future, we can expect further advances in this scientific field ([2]-Livezey and Glaser, 2020). Glaser et al (2020) attempted to analyze in his study entitled “Machine Learning for Neural Decoding”, with machine learning approaches, the decoding of spike activity. With data from subjects’ motor cortex, somatosensory cortex, and hippocampus, machine learning techniques such as support vector machines, neural networks, and boosted trees were used to correlate with external variables and make predictions. In this study, about 80% success was achieved with the Ensemble method in determining the instantaneous speed of movement from the sequence of spikes from the motor cortex. The accuracy was about 60% in the hippocampus ([14]-Glaser et al., 2020). Fang et al (2010), achieved 82% accuracy in decoding motion control with the Spiking Neural Network (SNN) approach ([15]-Fang et al., 2010). In their study entitled “Robust and accurate decoding of hand kinematics from entire spiking activity using deep learning”, Ahmadi et al. decoded hand kinematics from entire spiking activity obtained from the primary motor cortex of primates using the quasi-recurrent neural network (QRNN). The Pearson correlation coefficient was found to be about 90% in the classification of hand movements ([16]- Ahmadi et al., 2021).

The number of neural data decoding studies consisting of auditory stimuli with deep learning is increasing. In their study entitled “Study of Deep Learning for Sound Scale Decoding Technology from Human Brain Auditory Cortex”, Shigemoto et al. achieved 75% accuracy in classifying fMRI images of 3 subjects to which they applied different sound levels. He developed a sound scale decoding system for pitch using DBN and CNN ([17]- Shigemoto et al., 2019). Sridhar et al, in their study entitled “EEG and Deep Learning Based Brain Cognitive Function Classification”, achieved 65% success in 2020 using the BLSTM deep learning method on EEG signals obtained through auditory stimulation in an experiment involving 35 people ([18]- Sridhar et al, 2020). To reconstruct speech from human auditory cortex for brain-computer interface (BCI) solutions restoring communication for paralysis patients, Akbari et al. used the acoustic representation including auditory spectrogram and speech synthesis parameters with linear and non-linear regression methods. In this work, the DNN-vocoder combination achieved the best performance (75%accuracy)([19]- Akbari et al., 2019). Katthi et al, (2020) used methods such as canonical correlation analysis to analyse the relationships between the stimulus and EEG signals. They present a deep learning framework that learns to maximise correlation. The proposed method can decode human auditory attention ([20]-Katthi et al., 2020). Faisal et al, in their study, proposed a kernel convolution model to characterise neural responses to natural sounds. They used magnetoencephalography (MEG) on subjects receiving different sounds from the environment. By deducing the stimulus frequencies from neural responses, this model was able to precisely distinguish between two different sounds with 70% accuracy ([21]-Faisal et al., 2015). In their study entitled “Decoding Imagined and Spoken Phrases From Non-invasive Neural (MEG) Signals”, Dash et al. conducted 100 trials with 5 sentences that 8 people said or thought. They classified 3046 MEG signals using the ANN and CNN networks and obtained 96% accuracy for what was thought and 93% accuracy for what was spoken ([22]-Dash et al., 2020). Sharon et al, in their study entitled “Neural Speech Decoding During Audition, Imagination and Production”, achieved 54% success with the Gaussian mixture-based hidden Markov machine learning model from EEG signals collected from 30 individuals ([23]-Sharon et al.., 2020). Heelan et al, conducted two experiments in their study entitled “Decoding speech from spike-based neural population records in secondary auditory cortex of non-human primates” in 2019. 96 spike activities, obtained from 2 macaque monkeys hearing 30 words, were classified by Dense Neural Network (NN), Simple Recurrent NN (RNN), gated recurrent unit (GRU), long short-term memory (LSTM) RNN, having 0.54 successes in the ESTOI test. Furthermore, the average Mel-spectrogram correlation success rate with the machine learning methods, Dense Neural Network, simple RNN, GRU, LSTM was 90% in the classification of 20 English words spoken to 1 monkey ([24]-Heelan et al., 2019).

The dataset used in our study was prepared by Sadeghi et al. from their study “A neural ensemble correlation code for sound category identification”. They studied the correlation between the MUA signals measured in CI of two rabbits and five natural sounds that were played to them. Using spectrotemporal correlation for 1000ms in the acoustic paradigm1, he classified the MUA signals, based on the sounds heard, using statistical methods, and achieved a 90% accuracy value. In their work, Sadeghi et al. utilize a single-trial classifier and a noiseless classifier that takes the average of the validation data across trials, remove a major part of the variability across trials and isolate the stimulus-driven structure by increasing the accuracy. Both of them employ a Bayesian model to determine whether the five sounds can be distinguished by spectral or temporal neural correlation structure ([8]- Sadeghi et al., 2019).

A similar study, entitled “Distinct neural ensemble response statistics are associated with recognition and discrimination of natural sound textures” was conducted by Zhai et al. They investigated how natural sounds heard by 4 rabbits change the MUA signals measured in the CI in terms of correlation and spectrum. Using the neural spectrum method, they used a Naive Bayes classifier to classify the 1000 ms MUA signal segments according to the sounds heard. Taking the average of the results from the 29 different measurement sites, he obtained a maximum success rate of 90%. They obtained a higher accuracy value with shorter time windows compared to the results obtained by Sadeghi et al. In their work, Zhai et al. use natural and synthetic sound textures to prove that natural sounds’ statistical structure modulates the response statistics of the neural ensemble in the IC of unanesthetized rabbits (n = 4 animals, 29 penetration sites). The distinct statistics can potentially contribute to the recognition and discrimination of listened sounds. A naive Bayes classifier was trained with the neural correlations or neural spectrums of the original five sounds ([10]-Zhai et al., 2020).

The perceptual classification of a sound stabilises within about 1000 ms ([8]- Sadeghi et al., 2019, [10]- Zhai et al., 2020). For shorter durations, the classification success gradually degrades. We will try to increase these success values with image processing and deep learning methods. In the studies conducted, the success of the temporal correlation study was found to be lower. In our study, we will apply deep learning to temporal correlation images with processing windows of 250 ms and 1000 ms. Furthermore, we will see that the classification success increases significantly with the methods we use by removing noise with the cross-correlation technique on trials.

## II. MATERIAL AND METHODS

Environmental sounds, such as the crackling of fire, birdsong, crowds in the open air, running water and the sound of a snake, each includes unique time-frequency correlation characteristics. This allows us to verify and determine whether these features are encoded by the auditory midbrain. More specifically, we are interested in the temporal correlation representation in the activity of the neural ensemble.

### 2.1 DATA

The data are obtained from an international platform accessible at https://crcns.org/data-sets Collaborative Research in Computational Neuroscience ([25]- Dataset Paradigm 1-Sadeghi et al, 2019). In our study, with this dataset, 5 natural sounds will be predicted using the neural ensemble activity (MUA: MultiUnit Activity) signals from the inferior colliculus of two unanesthetized rabbits. In our study, we will try to find the heard sound stimulus using the Paradigm1 part of the dataset.

The neural data consists of multi-trial recordings of analog multi-unit activity (aMUA) measured from 16-channel linear recording arrays in the auditory midbrain (inferior colliculus) of unanesthetized rabbits receiving natural sounds. Paradigm 1 consist of five unaltered natural sound textures: fire, water, crowd noise, bird chorus, and rattling snake sound.

For each voice :

- 13 recording locations (sites): 4 from one animal, 9 from another,
- 16 channels for each recording site,
- 18-39 trials per channel.

Linear electrode array with multichannel acute neural recording silicon probes is used for neurophysiological recordings. Frequency response areas are shown for the recording channels ([10]-Zhai vd., 2020). For each of the 13 penetration sites, there are 16 channels (16 × 13 = 208 recording sites in the IC).

Instead of using the single unit activity which requires spike sorting or the thresholding multi Unit activity, it has been used an analog representation of multi-unit activity (analog multi-unit activity - aMUA) for several reasons ([8]-Sadeghi et al., 2019). Previous work indicates that aMUA signals include the structure of the population activity with a much lower noise level ([26]- Schnupp et al., 2015). According to Sadeghi et al., the measured neural signal is first bandpass filtered (325-3000 Hz), then full-wave rectified and low-pass filtered (475 Hz) ([27]-Rodriguez et al., 2010). These analog neural envelope signals, which are called aMUA, represent the synchronous activity and dynamics of the local neuronal population ([8]-Sadeghi et al., 2019).

### 2.2 TOOLS

#### 2.2.1 Correlation

The calculation of the cross-correlation coefficient makes it possible to account for the degree of association between two time series: this involves taking the synchronous values (x(t) and y(t)) in the two series. The cross-correlation coefficient can also be calculated by introducing a time lag between the two series: the calculation is then carried out by taking the pairs of values separated by a constant lag (x(t) and y(t+k)). These cross-correlation functions allow in particular the determination of the lag corresponding to the maximum association between the two variables. If a cross-correlation approach is applied to two non-linearly coupled series, taken as a whole, the shift corresponding to the maximum of the cross-correlation function will represent, as it were, the “average” of the shifts that could have been successively measured during the observation. One solution for analysing this type of phenomenon is to calculate the cross-correlation over a limited window, and then to slide this window along the analysed series ([28]-Delignières, 2007).

The autocorrelation of a series refers to the fact that in a time series, the measurement of a state at a time t can be correlated to previous measurements (at time t-1, t-2, t-3,…) or to subsequent measurements (at t+1, t+2, t+3,…). An autocorrelated series is thus correlated with itself, with a given time lag. Auto-correlation consists of correlating the series with itself, by introducing a lag between the two samples. The autocorrelation of lag 0 is equal to 1. The correlation decreases as the lag increases ([28]-Delignières, 2007).

As the data comes from the work of Sadeghi et al. ([25]- Dataset Paradigm 1-Sadeghi et al, 2019), based on their work entitled “A neural ensemble correlation code for sound category identification”, we can briefly say that, for each recording penetration, the neural ensemble correlations are calculated directly from the measured aMUA signals. The technique consists of a windowed short-term correlation modification in which the correlations are “ shuffled “ between the different response trials. The shuffling process is used to eliminate neural variation or noise from the recordings. To exclude the influence of response power on the total correlation measures, the short-term correlation is normalised as the correlation coefficient. ([8]-Sadeghi vd, 2019).

Temporal correlations are determined by computing autocorrelations between the same recording channels at different delays in the aMUA signals ([10]-Zhai vd., 2020). The calculations of temporal correlations require the detection of coincidences at different times. The temporal correlation structure of the aMUAs is shown by the **trial**-shuffled autocorrelograms (with the same channel) for each recording site ([8]-Sadeghi vd, 2019). The unshuffled auto-correlation calculation is with the same channel of the same trial.

#### 2.2.2 Preprocessing -Images

We have seen that the time-frequency correlations of sounds can be reflected by auditory assemblies in the midbrain. This will allow the identification and classification of natural sounds. Accordingly, the temporal correlations of the obtained aMUA signals are converted into images for different natural sound stimuli. The stimulus-induced temporal neural correlation images are used for classification. Thus, we will gain insight into the temporal structure and decoding of these neural signals.

Across sounds, the temporal correlations are very diverse and show a stimulus-dependent structure. For each sound, temporal correlations have a unique pattern and unique timescales. They display a brief peak at zero lag; however, they also exhibit a broader, slower correlation component that is stimulus dependent. For example, according to Sadeghi et al. “a relatively broad and slow component is observed for bird vocalization, and a periodic component at about 20 Hz is observed for rattlesnake noise.” ([8]-Sadeghi vd, 2019).

Neural correlations were calculated across the 16 recording channels. There are 5 natural sounds and 13 registration sites. The data used in this study consisted of two types :

- A **single trial “unshuffled”** is processed with autocorrelation. Here, noise is not removed. The channels are processed by autocorrelation. When it is a single trial, the single channel of that trial is auto correlated with the same channel. The unshuffled auto-correlation calculation is done with the same channel of the same trial. Two random windows are taken for each trial, so 2 x 5 x 13 x number of trials = 3180 images are generated.
- **“shuffled” noiseless**, cross-channel correlations across response trials, computed. According to Sadeghi et al., we can note : “By definition, noise correlations correspond to correlated firing rate fluctuations across repeated presentations of a stimulus that are unrelated to the sensory signal. Noise correlations are typically measured by correlating the firing rate residuals taken across repeated presentations of the stimulus”. By eliminating the trial-to-trial variability of neural activity, it was possible to remove the noise. Then the signal was separated into two parts: stimulus and noise. With this method we obtained noiseless images. For all trials, the temporal correlations are obtained by calculating the auto-correlations between the same recording channels with cross-correlation in trials. Example: channel 3 is correlated with channel 3, channel 12 is correlated with channel 12 in trial 3 and trial 4 selected by mixing. For each site, 50 images are randomly extracted, i.e. 5 x 50 x 13 = 3250 images. (This is for temporal correlation. If we had used spectral/spatial correlation, it would be both inter-trial and inter-channel).

Sound categorisation performance is highly dependent on the duration of the sound, as for human listeners (1-2 s). The relatively long time intervals, greater than 1 second, required for perceptual decision making might rather reflect an accumulation process. From a neural coding perspective, temporal resolution can be seen as the window of time in which neurons are computing correlations, while evidence accumulation time represents the time in which a neural population reads correlations to accumulate evidence. It is possible that sounds are integrated by the auditory system for decision making on relatively short time scales ([8]-Sadeghi vd, 2019). Rather than working only long time scales, a time window of 250 ms is also used. For both types of images, windows are taken with a duration of 250ms and 1000ms, corresponding to a ½ octave scale (Figure 3). During the acquisition of each window, the signal is clipped on both sides of the time series to eliminate artefacts. Maxium lag in ms for correlation is 100. (so the abscissas of the images are between −100 and +100)

**Figure 3:**
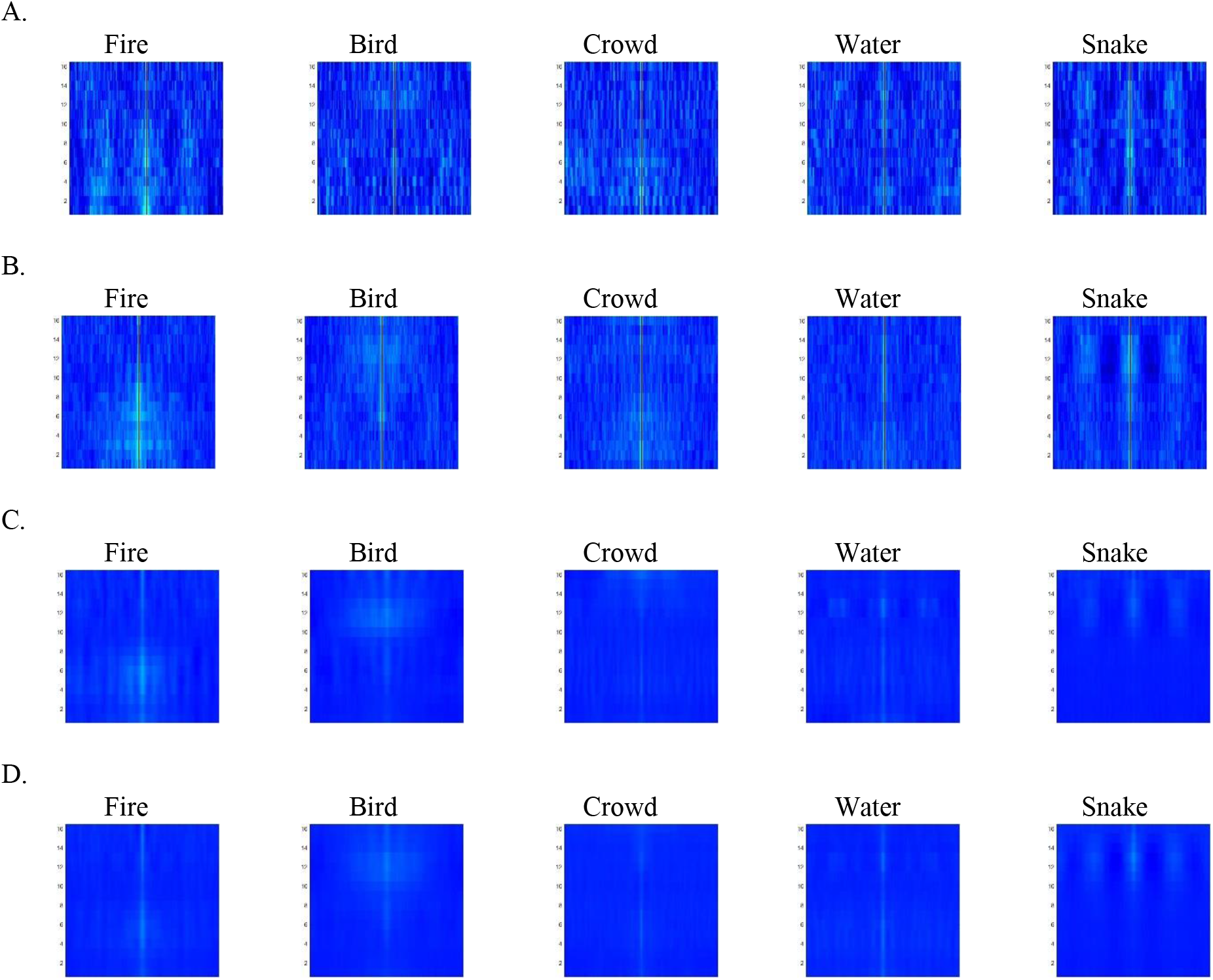
Time correlation images obtained for each sound. A. For a single trial and time windows of 250 ms. B. For a single trial and 1000 ms time windows. C. Noiseless and time windows of 250 ms. D. Noiseless and 1000 ms time windows.

The classifier identifies the delivered sound using single-trial response and all trials (noiseless) with windows of two different durations (250 and 1000 ms).

In the absence of cross-validation (in the case of hyperparameter optimization), the images are used at 0.7 for training and 0.3 for test. With cross-validation, the value of kfold being 10, the data are used at 0.9 for training and 0.1 for testing and the average acurracy value is calculated (Figure 5).

**Figure 4:**
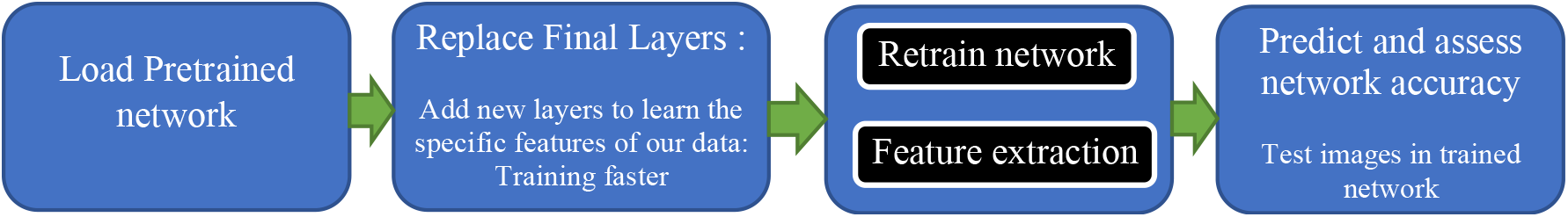
Reuse Pretrained Network.

**Figure 5.**
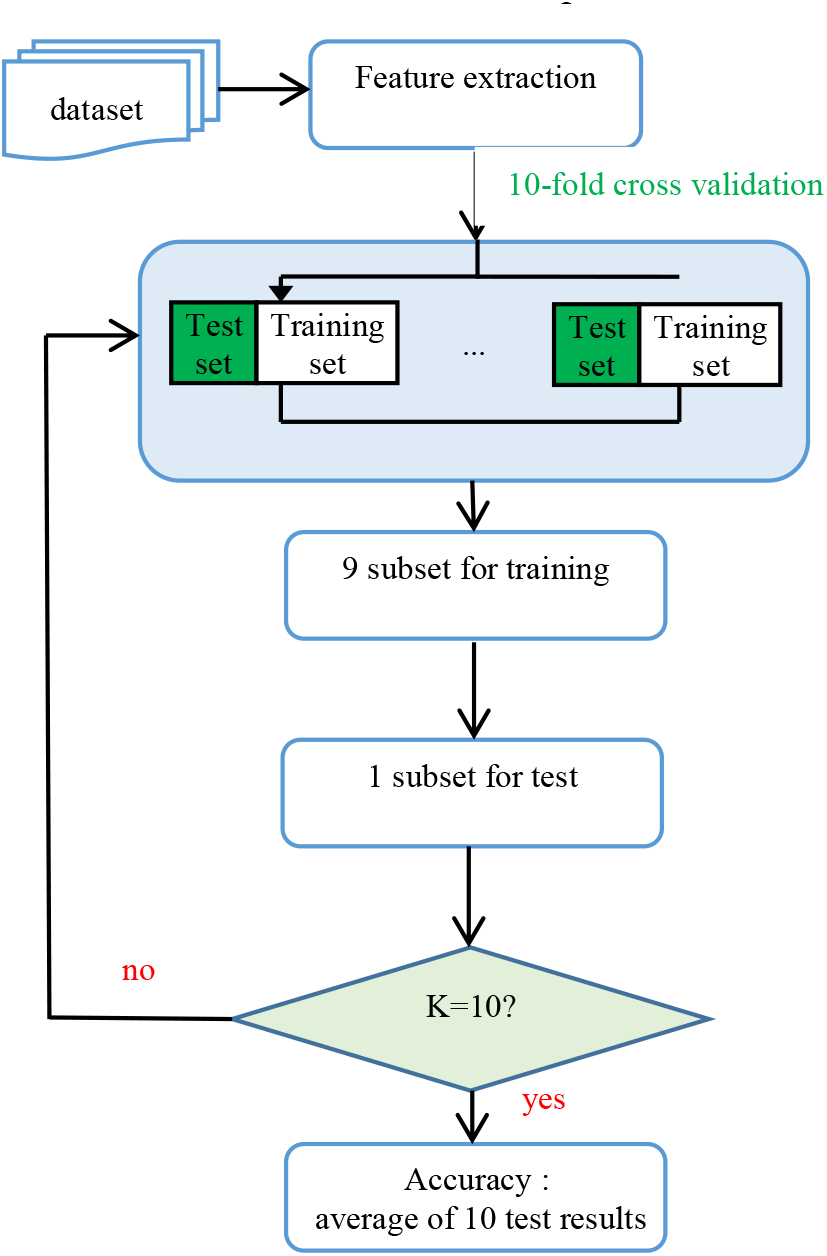
Classification with 10 cross-validation

Data reduction is one of the most frequently used pre-processing methods. Reducing the data size allows learning algorithms to work faster and more efficiently. The images are RGB, 227×227×3 in size and in jpeg format. In our study, feature selection was performed automatically by convolutional neural networks (CNN). As there is no large dataset, data augmentation and feature extraction techniques with transfer learning were used.

#### 2.2.3 Process

In this study, it is shown how to incorporate pre-trained deep networks to predict natural sounds from neural data. For this purpose, 4 types of images were used.

In order to obtain fast and efficient results, processing and classification were performed using 2 methods:

- Fixed decoder: Feature extraction with transfer learning and classification with 10 kfold cross validation.
- Retrained decoder : retraining the pretrained network (Alexnet) and classification with 2 kfold cross validation.

Accuracy, specificity, sensitivity, precision and F1 score metric values were calculated as parameters to evaluate the results. Finally, the values obtained with the best performing methods were compared.

##### a) Deep learning

It is possible to achieve much greater success in this study using deep learning techniques that have proven successful in image processing. In this work, pre-trained CNNs (Convolutional Neural Networks) such as AlexNet, GoogleNet and SqueezeNet was used to achieve the best results in terms of accuracy performance and computation time. CNNs, like multilayer perceptrons, have a lower number of parameters than other fully connected networks and are more easily trained ([29]-Özcan and Alkan, 2021).

It is usually much quicker and easier to use a pre-trained network with transfer learning than to train a new network. It is important to choose the right pre-trained networks to apply to our problem. For this, the choice is based on the different most important characteristics such as accuracy, speed and size of the network. The choice of a network is usually a compromise between these characteristics. We use pre-trained image classification networks that have previously learned to extract meaningful and relevant features from about one million images. These networks can classify images into 1000 different categories. As shown in Table 1, three pre-trained networks with different depth, number of layers, number of parameters and network size were selected ([30]- Matlab 2020b).

**Table 1.** Three available pretrained networks and their properties

| Network | Depth | Size | Parameters (millions) | Image input size | Number of layers | Number of connections |
| --- | --- | --- | --- | --- | --- | --- |
| Alexnet | 8 | 227MB | 61,0 | 227-by-227 | 25 | 24 |
| Googlenet | 22 | 27MB | 7,0 | 224-by-224 | 144 | 170 |
| Squeezenet | 18 | 5,2MB | 1,24 | 227-by-227 | 68 | 75 |

The Alexnet network is a convolutional neural network with 8 layers of depth. The use of an augmented image database allows us to get real results, as it prevents the network from over-adapting and memorising the exact details of the training images. Googlenet is a convolutional neural network that is 22 layers deep. Finally, Squeezenet is a convolutional neural network that is 18 layers deep. Table 1 lists the available pre-trained networks trained on ImageNet and their properties. The depth of the array is defined as the number of convolutional or fully connected sequential layers from input to output. RGB images can be presented at the input of the networks listed below ([30]-Matlab 2020b).

In transfer learning, to re-train a pre-trained network to classify new images, the last layers are replaced by new layers adapted to the new dataset. It is necessary to modify the number of classes according to the latter (here 5). Thus, the network classifies the input images using the new last layers (training and final classification) ([31]-L. Iliadis et al.). The last trainable layer of the network is a fully connected layer.

##### b) Fixed decoder : Feature extraction

From a pre-trained convolutional neural network, we will extract previously learned image features, and use them to train our image classifier. Feature extraction is a quick and easy way to use the power of deep learning without investing time and effort in training a complete new network. If a GPU is not available, it is particularly useful as it only requires a single pass over the training images. It is therefore possible to adapt a multiclass classifier and classify the test images. The feature extraction solution should be chosen when the new data set is small. The network establishes a hierarchical representation of the input images. The deeper layers contain higher level features, built from the lower level features of the preceding layers. To obtain the feature representations of images, it is necessary to use the activations of the fully connected layer ([30]-Matlab 2020b).

With the feature extraction method, the image features were extracted and the type of sound heard was classified. The Alexnet Googlenet and Squeezenet CNNs were used for this purpose.

##### c) Retrained decoder : Transfer learning and retrain neural network

In deep learning applications, transfer learning is frequently used. By using a smaller number of images for the training process, we can quickly transfer previously learned features to a new task. As a starting point, we can fine-tune the deep layers of the pre-trained network by re-training the network on our new data set. Fine-tuning an existing, pre-trained network is often faster and easier than building and training a new one. The network having already learned a rich set of image features, can therefore learn features specific to our dataset. Fine-tuning a network is slower and requires more effort than simple feature extraction but often gives the best accuracy ([30]- Matlab 2020b). The Alexnet CNN, which is first trained with a large dataset, can be refined on our neural data.

We can see in the following Figure 4 the method of reusing a pretrained network.

##### d) Classification and evaluation methods

The principle of image classification consists in classifying images into different categories, according to a label. After feature extraction for the fixed neural decoder, we used 4 different classifiers methods to compare results: Support Vector Machine (SVM), K-Nearest Neighbor (kNN), Naive Bayes, Ensemble. SVM, Naive Bayes and Ensemble classifiers have been briefly explained in our previous work ([29]-Özcan and Alkan, 2021). k-nearest-neighbor (KNN) classification model allows us alter both the distance metric and the number of nearest neighbors.

Selecting the value of hyper-parameters plays a crucial role in classification. In order to determine the most adequate hyperparameters for our classifications, we have done an optimization work. In all classification algorithms, the hyperparameter optimization option is used. For this, 30 tests are performed. By selecting the optimal value of the hyperparameters, the best performance of the classifier can be obtained. As the processing time for hyperparameters optimization, we did not apply a cross-validation, and the data were splited into 70% for training and 30% for test.

One of the validation techniques used to test the accuracy of the model when the number of data is limited is the K-Fold cross-validation method. The cross-validation method is the most accurate approach to training and classifying data. It divides the data set into K groups of equal size (here, 10). One group is always reserved for test, while the trained model is built with the other groups. The accuracy rate is calculated with the data allocated for testing. This situation is repeated iteratively K times. Thus, each group is used the same number of times for training and test. The accuracy rate of the model is the average of the K accuracy rates ([32]-Fakı, 2015). In experiments that did not require a huge amount of time for processing, a 10-fold cross-validation was applied (Figure 5). In this work, a 10-fold cross-validation technique was used.

Accuracy, is calculated to compare each result according to pre-trained neural network and classification methods. Specificity, Sensibility, Precision, Recall, F1-Score were calculated for the case that feature extraction with Alexnet and classification with SVM and hyperparameter optimization for the fixed decoder applied on 250ms-single trial, 1000ms-single trial, 250ms-noiseless and 1000ms-noiseless images.

Confusion matrices were generated for each model. It is used in the classification results to evaluate the accuracy of the model. The confusion matrix allows us to detect the areas where the classifier is less efficient. In the table, we can see the actual classes and the predicted classes in rows and columns. The diagonal cells indicate the degree of correspondence between the actual class and the predicted class.

ROC curves were generated for the re-trained decoder applied to 250 ms single trial, 1000 ms single trial, 250 ms noiseless and 1000 ms noiseless images. The Receiver Operating Curve (ROC) is one of the widely used methods for assessing the accuracy and reliability of models. Each point on the ROC curve corresponds to a model created by the classifier by choosing a fixed threshold (Figure 12B). The ROC curve classifies the units into classes using the sensitivity and specificity ratios and determines the most appropriate cut-off point. On the coordinate plane of the ROC curve, the X-axis represents the false positive rate (1-specificity) and the Y-axis the true positive rate (sensitivity) ([32]-Faki, 2015). A perfect result is represented by a right angle in the upper left corner of the area. A poor result is shown as a straight line at 45 degrees. The overall quality of the classifier is assessed by the area under the curve (AUC). The AUC value indicates the separability of the groups to be classified. This value is independent of the classification threshold. Higher AUC values indicate better performance of the classifier. An AUC of 1 means that 100% of the class groups are separated (AUC ≥ 0.9, excellent; 0.8 - 0.89, good; 0.7 - 0.79, borderline; others, unacceptable). If a selected classifier model fails to classify correctly and select the threshold value correctly, the AUC may be high and the success low.

**Figure 6:**
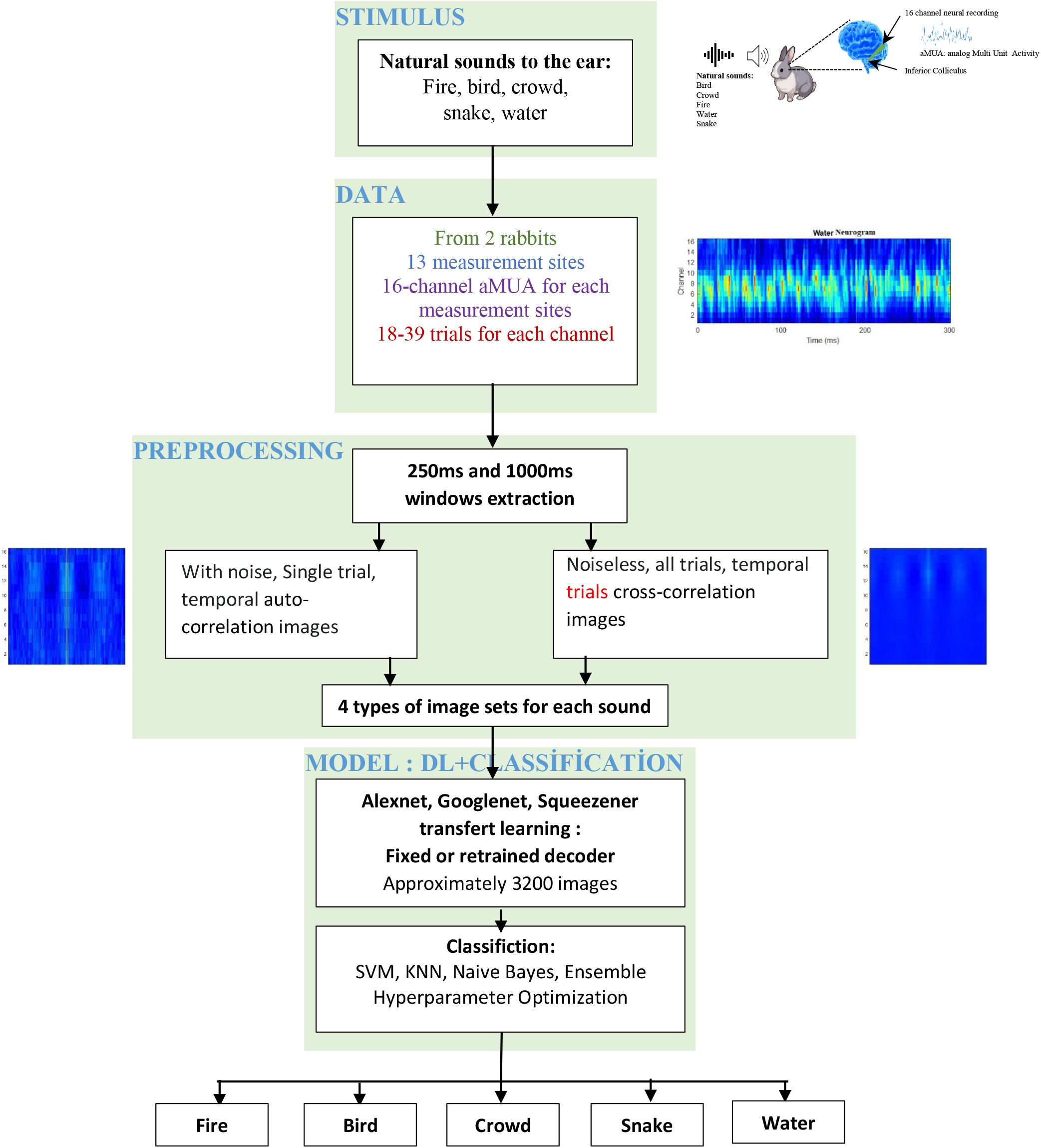
Flow chart of the study, from IC measurements to the classification of the sound stimulus.

**Figure 7.**
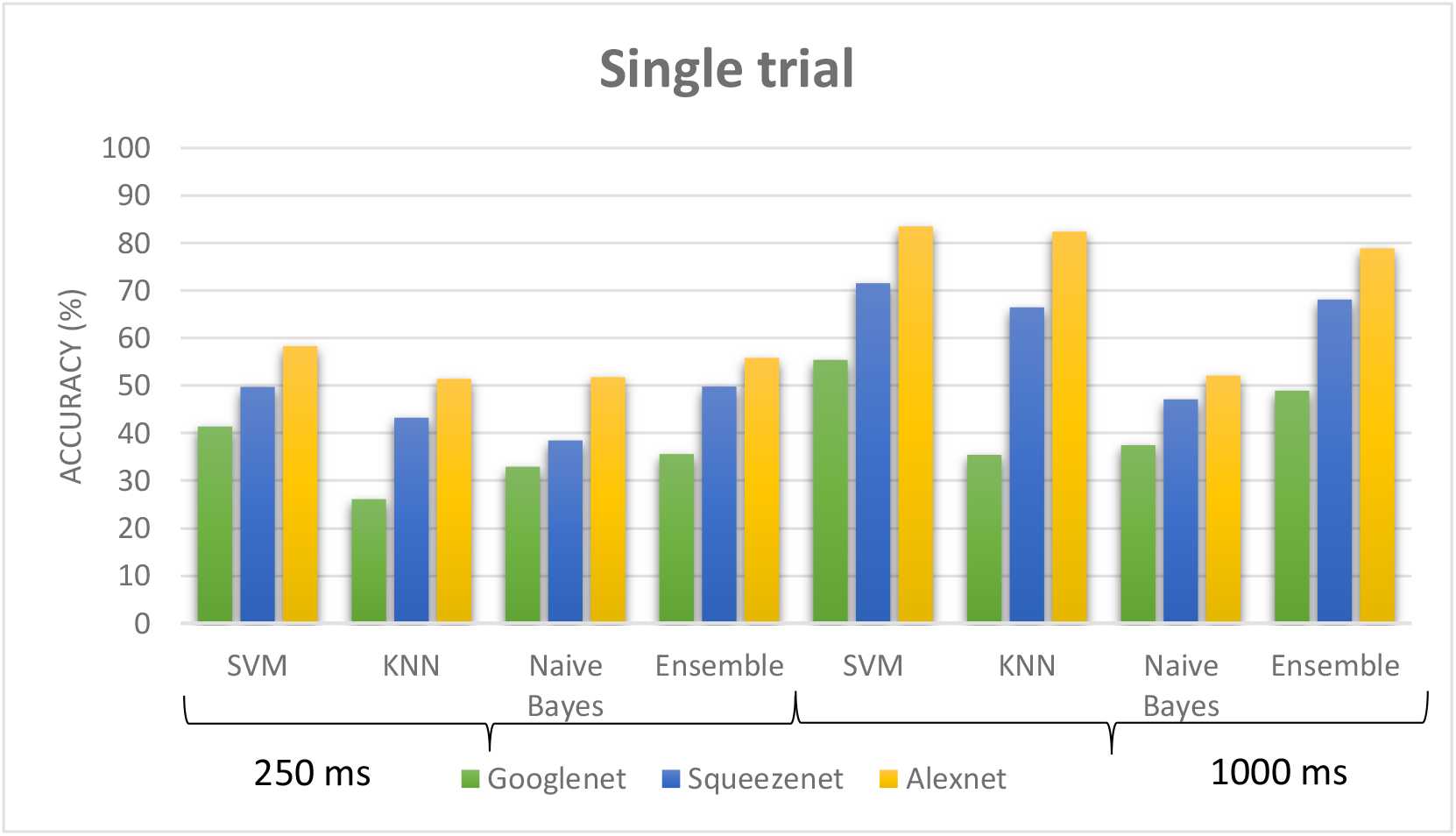
Decoding performance (accuracy) comparison across different window size, different network and different classifiers for single trial processing.

**Figure 8.**
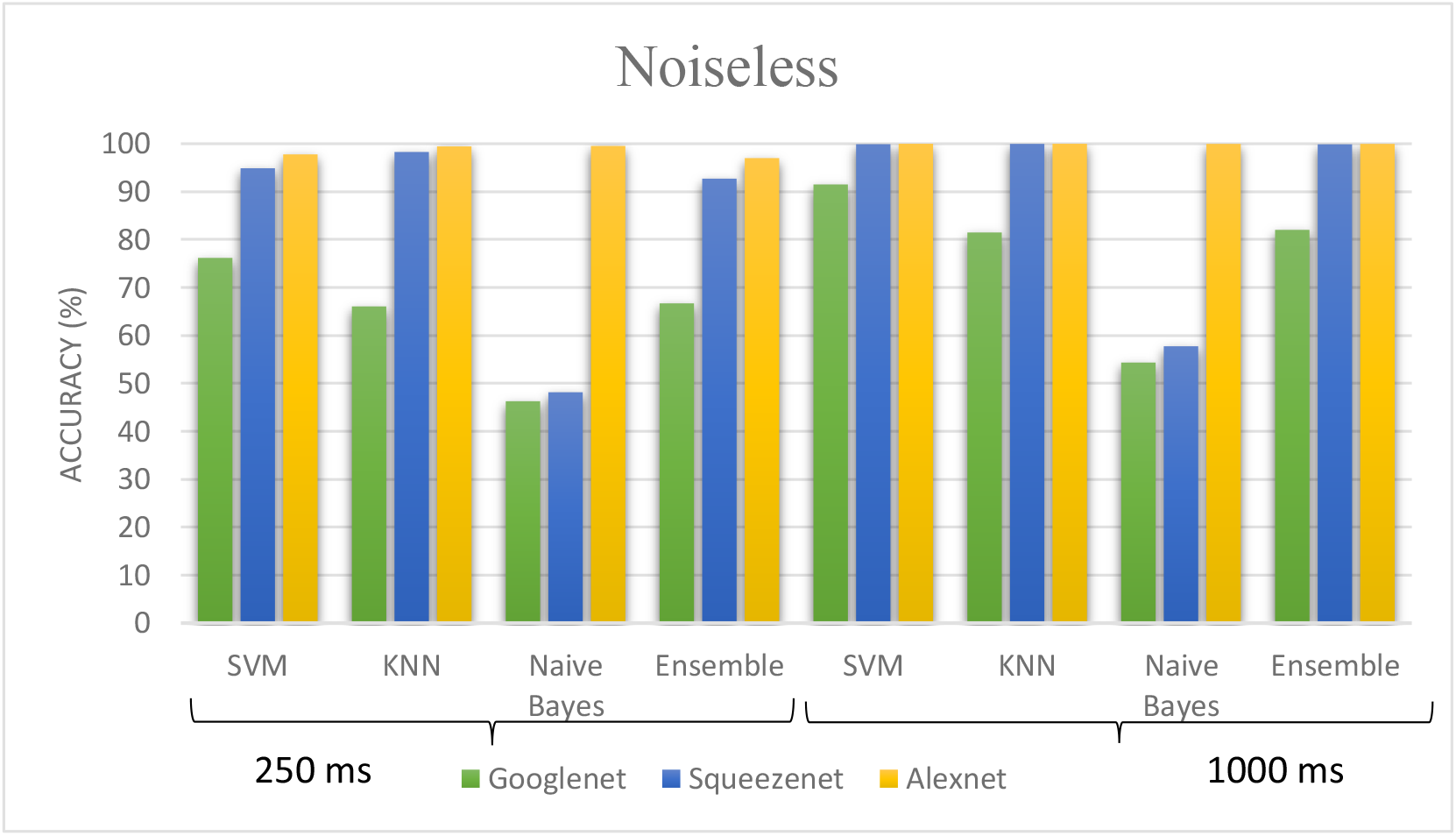
Decoding performance (accuracy) comparison across different window size, different network and different classifiers for noiseless processing.

**Figure 9.**
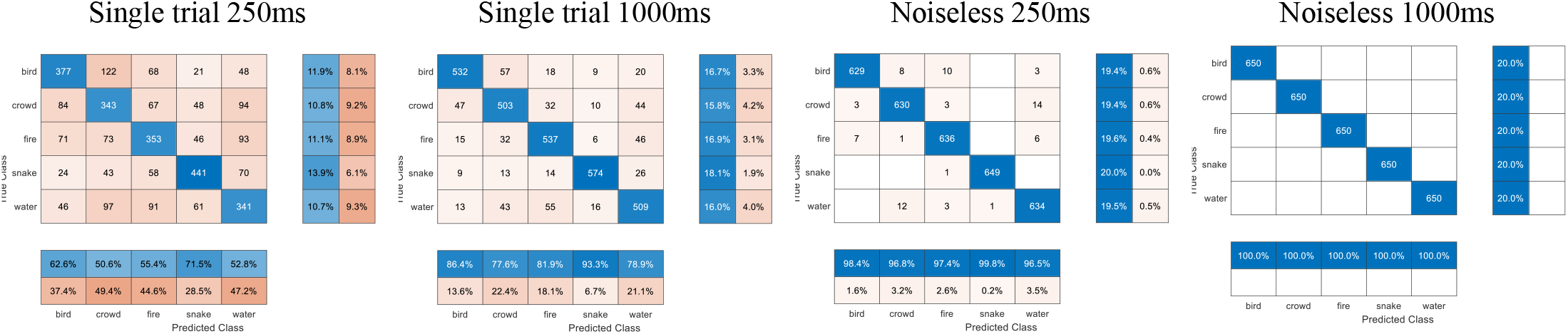
Decoding performance (accuracy) comparison across different classes with Alexnet transfer learning and SVM classifier.

**Figure 10.**
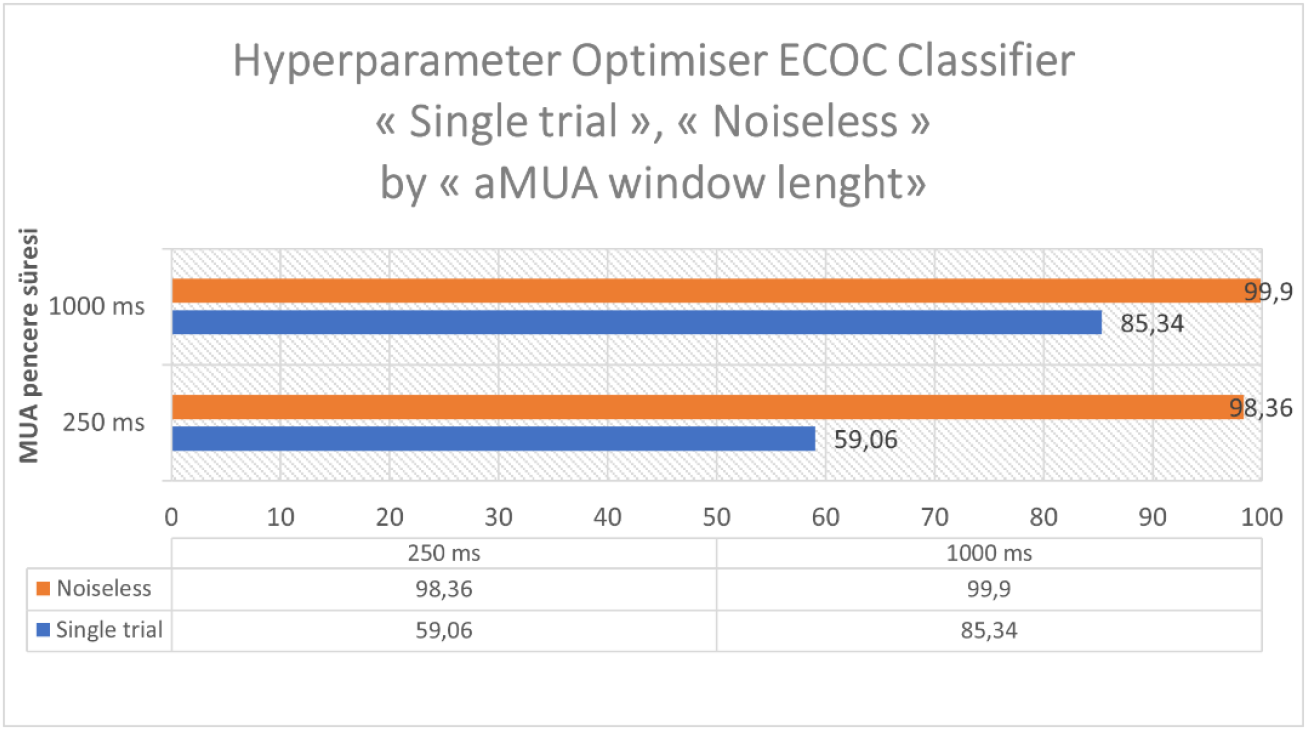
Hyperparameter Optimization for ECOC classifier: accuracy performance according to window duration.

**Figure 11.**
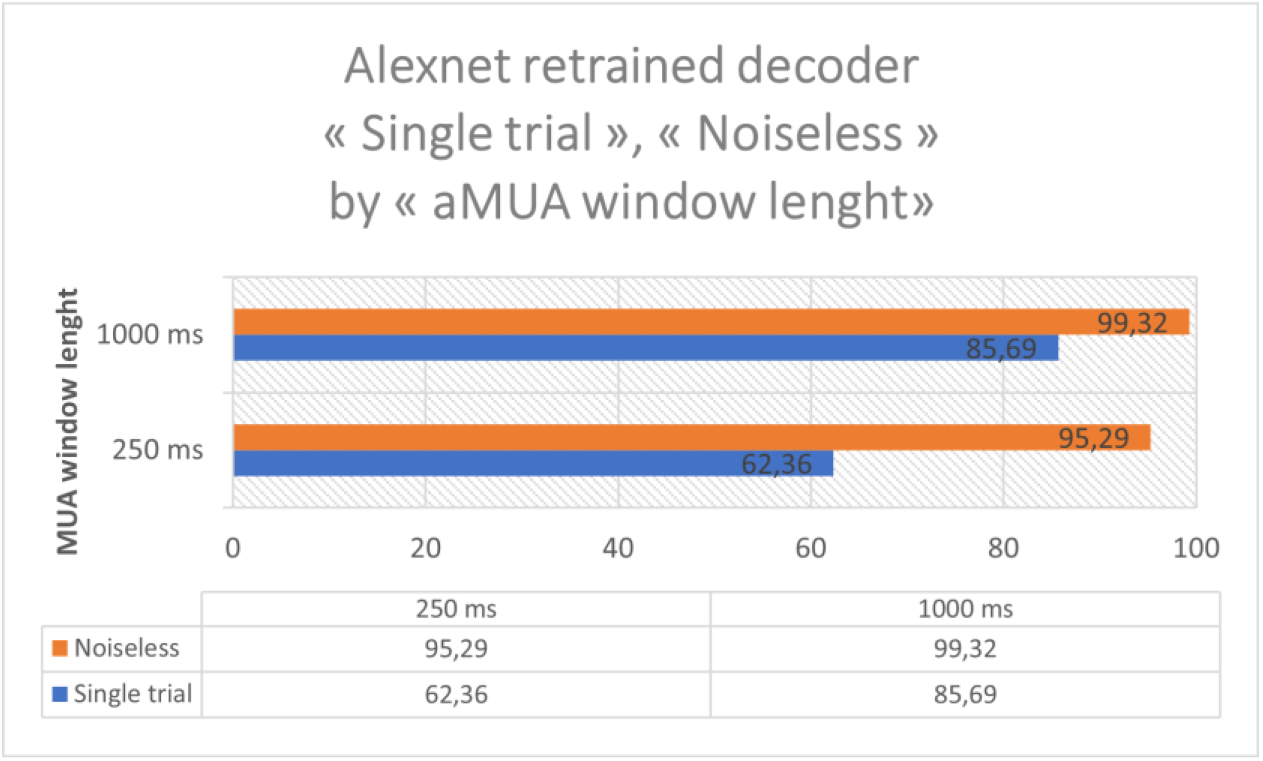
Retrained decoder accuracy performance according to windows duration

**Figure 12.**
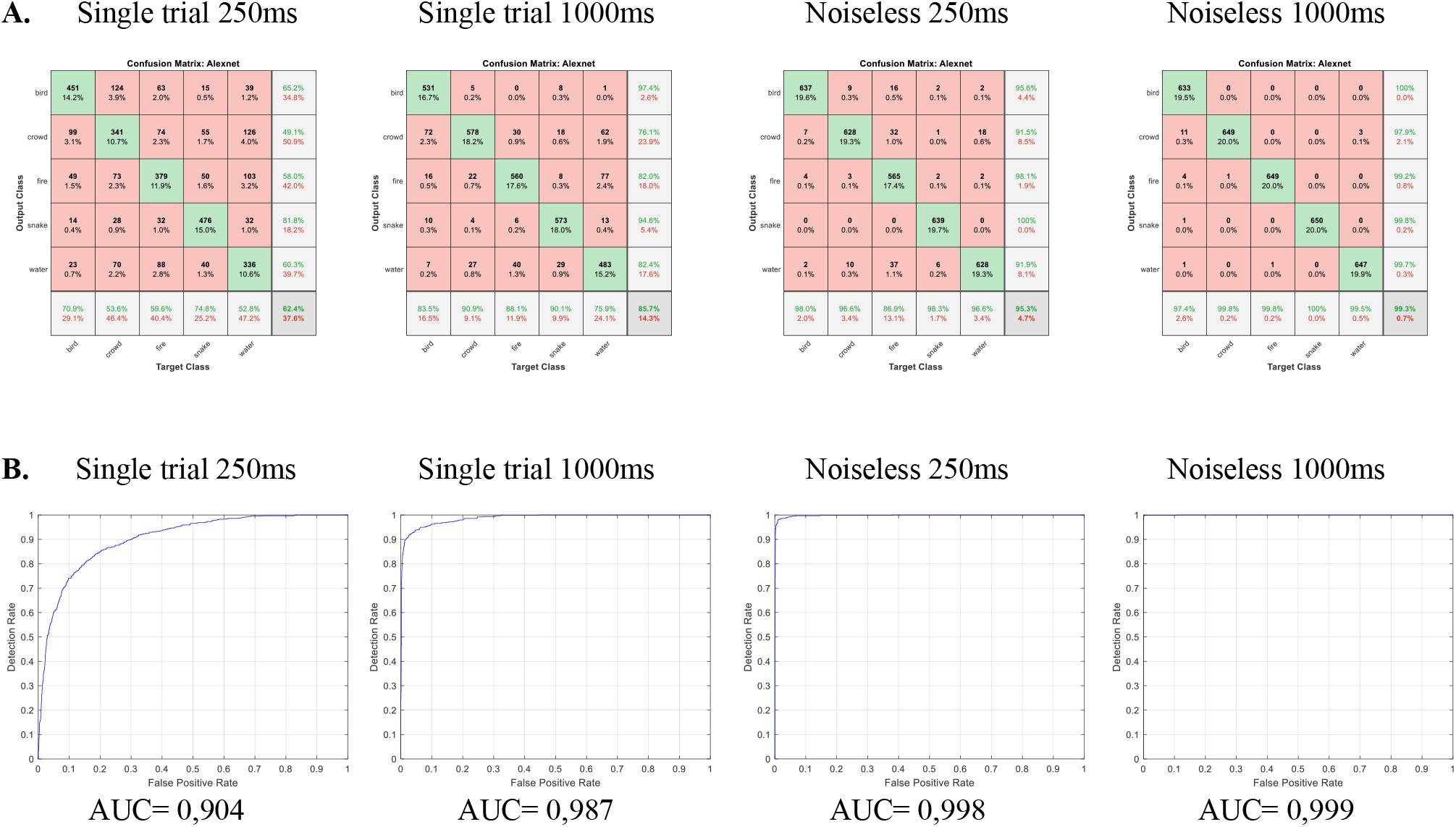
Retrained decoder accuracy performance (A) and ROC curve (B) according to window duration across different classes with Alexnet transfer learning.

## III. RESULTS AND DISCUSSION

### 3.1 RESULTS

In this work, the classifier had to identify five sounds based on neural activity (Figure 6).

The classifier identifies the delivered sound using single-trial and noiseless responses with windows of two different durations (250 ms and 1000 ms). The cross-validated data were used to train and test the model. The classification performance is presented in tables 2 and 3.

**Table 2.**
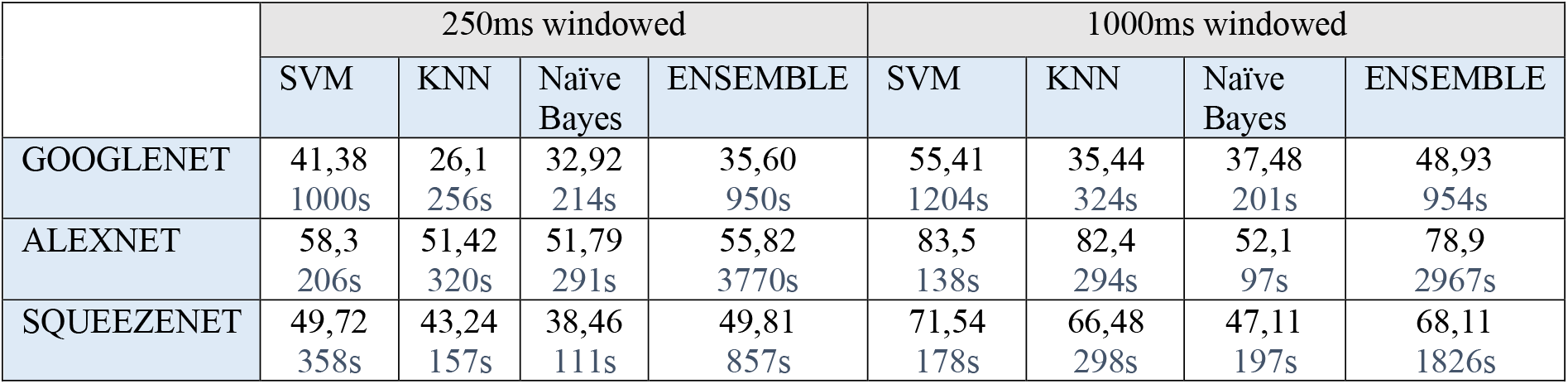
Single trial fixed decoder accuracy performance and processing time

**Table 3.** Noiseless fixed decoder accuracy performance and processing time.

|  | 250ms windowed |  |  |  | 1000ms windowed |  |  |  |
| --- | --- | --- | --- | --- | --- | --- | --- | --- |
|  | SVM | KNN | Naïve Bayes | ENSEMBLE | SVM | KNN | Naïve Bayes | ENSEMBLE |
| GOOGLENET | 76,18<br>642s | 66,06<br>529s | 46,28<br>301s | 66,7<br>1763s | 91,51<br>223s | 81,48<br>256s | 54,34<br>207s | 82,03<br>1005s |
| ALEXNET | 97,78<br>111s | 99,45<br>308s | 99,54<br>316s | 97<br>3596s | 1<br>101s | 1<br>303s | 99,97<br>99s | 99,97<br>3803s |
| SQUEEZENET | 94,89<br>153s | 98,28<br>159s | 48,15<br>107s | 92,71<br>871s | 99,91<br>111s | 99,97<br>157s | 57,75<br>104s | 99,88<br>881s |

We will do decoding performance comparison across different cases (single trial, noiseless), different window size, different fixed and retrained networks and different classifiers.

#### 3.1.1 Fixed decoder : Feature extraction with transfer learning and classification

The fixed decoder consists in using transfer learning with pretrained network for feature extraction in order to classify the data according to the case (single trial, noiseless), the window duration, the different networks and the different classifiers. 10 kfold cross validation is applied for classification.

The highest success value we obtained in classifying the 250 ms windows recorded with a single trial was 58.3% (Table 2 and Figure 7). This value was obtained with the SVM classifier by Alexnet feature exctraction. Determining the parameter values of the classifier with hyperparameter optimization gave 59.06% (Figure 10). By re-training the Alexnet with our own data, we obtained the best result, 62.36% (Figure 11). In this case, as shown in Figure 12, the highest classification success was obtained for the snake class with 74.8% and for the bird class with 70.9%, and the lowest was obtained for the water class with 52.8%. The sensitivity of the crowd class (53.6%) was relatively low.

The highest accuracy value we obtained in the classification of 1000 ms windows recorded with single trial was 83.5%. This value was obtained with the SVM classifier by Alexnet feature extraction. Squeezenet, one of the other networks we used, gave better results than Googlenet. These results were obtained with cross validation (kfold=10). By determining the value of the classifier parameters with hyperparameter optimization, 85.34% was obtained (Figure 10). Compared with SVM, KNN and Naive Bayes classifiers, processing time is relatively long for Ensemble classifier. By retraining the Alexnet network with our own data (fine-tune), the highest accuracy was obtained as 85.69% (figure 11). In this case, as can be seen in Figure 12, the classification accuracy was highest for the crowd class with 90.9% and for the snake class with 90.1% and lowest for the water class with 75.9%.

According to the success values, we can order the networks as Googlenet < Squeezenet < Alexnet and the classifiers as Naive Bayes < KNN < Ensemble < SVM. With high performance, we can see the shortest process time for Alexnet and SVM.

After noise removal, using all trials, the highest accuracy rate for classification of the recorded 250 ms windows was 99.54% (Table 3 and Figure 8). This value was obtained with the Naive Bayes classifier by feature extraction with the Alexnet network. When determining the value of the SVM parameters with hyperparameter optimization, the accuracy was 98.36% (Figure 10). By re-training the Alexnet with our own data, we achieved 95.29% success (Figure 11).

In noiseless case, the highest accuracy value we obtained in the classification of the recorded 1000 ms windows was 100%. This value was obtained with the SVM and KNN classifiers by feature extraction with the Alexnet network and cross validation (kfold=10). With hyperparameter optimization, the classifier score was 99.9%. Less success was achieved by re-training the Alexnet with our own data, with an accuracy of 99.32%.

We can see confusion matrix for each case in Figure 9.

With the hyperparameter optimization on the ECOC classifier, as we can see in Figure 10, we obtain better results on single trial classification. We don’t use cross validation in hyperparameter classification because the process time is long (about one hour).

The noiseless process allows for a significant increase in classification results, even with short time windows.

In Table 4, 5, 6, 7, we can see different evaluation value across classes with hyperparameter optimization in an ECOC classification.

**Table 4.** Single trial – 250 ms

HYPERPARAMETER OPTIMIZATION on ECOC Classifier results
|  | Specificity | Recall/Sensitivity | Precision | F1 score |
| --- | --- | --- | --- | --- |
| Bird | 93,46 | 58 | 68,9 | 62,98 |
| Crowd | 83,12 | 58,5 | 46,5 | 51,81 |
| Fire | 88,61 | 57,5 | 55,8 | 56,64 |
| Snake | 93,85 | 69,5 | 73,9 | 71,63 |
| water | 89,8 | 51,5 | 55,7 | 53,52 |
| Average - Total | 89,768 | 59 | 60,16 | 59,32 |

**Table 5.**
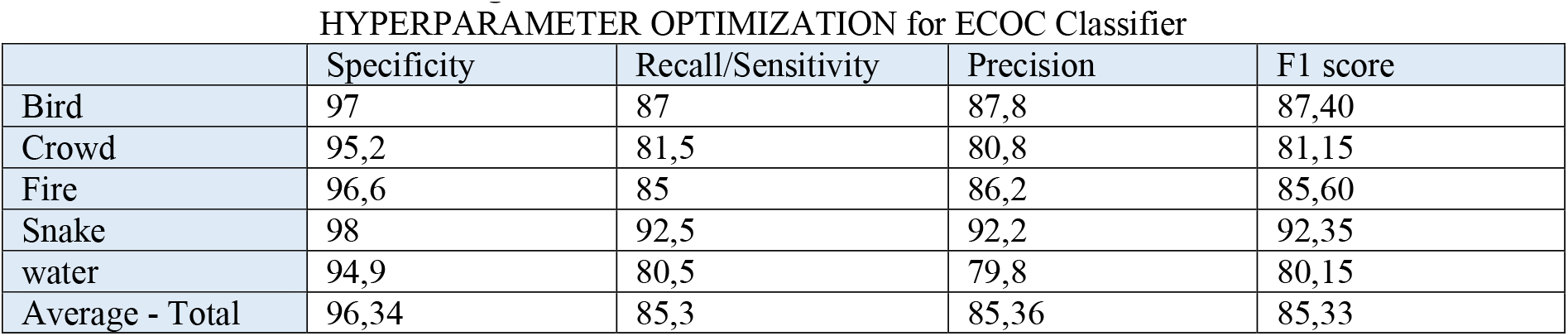
Single trial – 1000 ms

HYPERPARAMETER OPTIMIZATION for ECOC Classifier
|  | Specificity | Recall/Sensitivity | Precision | F1 score |
| --- | --- | --- | --- | --- |
| Bird | 97 | 87 | 87,8 | 87,40 |
| Crowd | 95,2 | 81,5 | 80,8 | 81,15 |
| Fire | 96,6 | 85 | 86,2 | 85,60 |
| Snake | 98 | 92,5 | 92,2 | 92,35 |
| water | 94,9 | 80,5 | 79,8 | 80,15 |
| Average - Total | 96,34 | 85,3 | 85,36 | 85,33 |

**Table 6.** Noiseless – 250 ms

HYPERPARAMETER OPTIMIZATION for ECOC Classifier
|  | Specificity | Recall/Sensitivity | Precision | F1 score |
| --- | --- | --- | --- | --- |
| Bird | 99,49 | 96,5 | 97,9 | 97,19 |
| Crowd | 99,49 | 98,5 | 98 | 98,25 |
| Fire | 99,62 | 98 | 98,5 | 98,25 |
| Snake | 100 | 100 | 100 | 100,00 |
| water | 99,36 | 99 | 97,5 | 98,24 |
| Average - Total | 99,592 | 98,4 | 98,38 | 98,39 |

**Table 7.** Noiseless – 1000 ms

HYPERPARAMETER OPTIMIZATION for ECOC Classifier
|  | Specificity | Recall/Sensitivity | Precision | F1 score |
| --- | --- | --- | --- | --- |
| Bird | 100 | 100 | 100 | 100,00 |
| Crowd | 99,87 | 100 | 99,5 | 99,75 |
| Fire | 100 | 99,5 | 100 | 99,75 |
| Snake | 100 | 100 | 100 | 100,00 |
| water | 100 | 100 | 100 | 100,00 |
| Average - Total | 99,974 | 99,9 | 99,9 | 99,90 |

According to the Table 4, by applying feature extraction and hyperparameter optimization with the Alexnet network, the highest sensitivity we obtained in classifying the 250ms windows recorded with a single trial was for the snake (69.5%) and crowd (58.5%) classes. The highest specificity and precision values were obtained for snake and bird classes. If we consider the metric value of the F1 score, we can say that the classifier performs well in the snake and bird class. The lowest F1 score value was obtained with crowd class.

According to the Table 5, by applying feature extraction and hyperparameter optimisation with the Alexnet network, the highest sensitivity we achieved in classifying 1000ms windows recorded with single trial was for bird and snake classes. In addition, the highest specificity, sensitivity, precision and F1 score values were obtained for these two classes. The lowest values were obtained with water and crowd.

By extracting features with the Alexnet network, using the HO classifier, the highest success we achieved in classifying the 250ms noiseless windows was for the snake (100%) class. The highest specificity, sensitivity, precision and F1 score values were obtained for this class. The other classes have almost the same values. The lowest sensitivity and F1 score values were obtained with the bird class. According to the Table 6, with noiseless images, we can say that the classifier performs well in the snake, crowd, fire and water class.

According to the Table 7, with noiseless images, we can say that the classifier performs well in all classes.

#### 3.1.2 Retraining decoder : Alexnet Retrained decoder, 2 cross validation, and classification

The retrained decoder consists in using pretrained network, Alexnet, for retrain network with our data in order to classify according to the case (single trial, noiseless) and the window duration. 2 kfold cross validation is applied for classification. The process duration is about one hour for each retraining act.

According to the results on figures 15 and 16, we obtain similar results comparing to the fixed decoder.

### 3.2 DISCUSSION

Two unanesthetised rabbits were stimulated with five natural sounds and multiunit, multichannel neural recordings were obtained from the auditory midbrain (inferior colliculus) to extract aMUAs. The aMUA signals were cut into 250 ms and 1000 ms windows. These signals were classified for the decoding of the neural data. Thus, it was shown that our model was also efficient for decoding short time integration. All the work we have done has focused on temporal correlation. The accuracy of the classification was improved by eliminating, during pre-processing, the noise in the data with the cross-correlation method across trials. We verified that natural sounds affect the correlation structure in the neural data, and that this in turn affects the recognition of the sound category.

In a study with the same data, Sadeghi et al. achieved a success rate of about 50% with 250 ms windows and 65% with 1000 ms windows in classifying temporal correlation with single trial and data from 13 sites. If we consider the difference in the highest accuracy value (85.69%) that we obtained for the 1000 ms images, we see an increase of 31.83%. Sadhegi et al. obtained about 83% with spectro-temporal correlation (single trial, 1000ms, average of 13 sites). In our study, we obtain more than this value just using the temporal correlation. If we consider the difference of the highest accuracy value we obtained for 250ms (62.36%), we see an increase of 24.72% for the temporal correlation. In this scheme, we find that the classification of fire and water with a single site was low (about 30%) for Sadhegi et al. In our average of 13 sites, this classification increased to 100%. In the study by Sadeghi et al, fire and water were the classes classified unsuccessfully with single site data in the temporal correlation classification ([8]- Sadeghi et al, 2019). The best classified classes were snake, bird and crowd. These results are consistent with our results for the snake and bird classes.

Again, in the Sadeghi et al. study, in the noiseless case, an average of about 65% success was obtained in temporal correlation classification with data from 13 sites and 1000ms windows. If we consider the highest success value difference we obtained (100%), we saw that our accuracy increased by 53.85%. In this case, in spectro-temporal correlation, Sadhegi et al. obtained an average of about 90%. As we obtained a 100% success rate with temporal correlation in our study, we can see that the method we used is much more successful. On the other hand, in the Noiseless case, an average of about 72% accuracy was obtained in temporal correlation classification with data from 13 sites and 250ms windows. As we obtained more than 99% in our study, we see an increase of 37.5%.

With similar data, Zhai et al, in their study entitled “Distinct neural ensemble response statistics are associated with recognition and discrimination of natural sound textures”, achieved about 90% success with 1000ms windows and about 78% success with 250ms windows with the method called Neural spectrum ([10] - Zhai et al., 2020). For 1000ms, our accuracy was 10% higher, for 250ms, our accuracy has a difference of 27.62%.

With the temporal correlation images, we obtained the highest value with 1000ms aMUA windows and reached the target.

In general, for all sounds, the performance of the classifier increases with the duration of the sound. For fire and water sounds, we can register a clear increase in accuracy with both types of windows.

With Alexnet, the classification accuracy is indeed much better than other CNNs. In addition, compared to advanced CNNs such as GoogLeNet or SqueezeNet, AlexNet consumes less time in the classification process. According to the success values, we can rank the networks as Googlenet < Squeezenet < Alexnet and the classifiers as Naive Bayes < KNN < Ensemble < SVM. Together with high performance, we can see the shortest process time for Alexnet and SVM. The fixed decoder gives faster results than the retrained decoder and the success results are approximately the same.

Here we can emphasise that the squeezenet network is very close to Alexnet, although the number of parameters is small. Alexnet and Squeezenet performed better than Googlenet.

Sound categorisation performance is highly dependent on the duration of the sound, as for human listeners (1-2s). It is possible that sounds are integrated by the auditory system for decision making on relatively short time scales. Rather than working on long time scales, a time window of 250 ms is used also, which also corresponds to a 1/2 octave value ([8]-Sadeghi et al. matla, 2019). Considering that humans hear and classify sounds in about 1 second, a high segment classification success of 250 ms gives us hope for future human-machine interface designs. All the work we have done has focused on temporal correlation and. Temporal correlation appears to be very effective in decoding neural data. The tonotopic order of the temporal correlation provides information. It was verified that noise limits the encoding of sensory information. Since the noise in the data was eliminated by the cross-correlation method, the success of the classification was improved. The performance of the noiseless images classification is superior to that of the single-trial images classification. The AUC value results for 250 ms windowed single trial and noiseless data, are very high and encourages us to work with shorter windows.

Here we have shown that natural sounds can induce temporal correlations between neural reponses from the inferior colliculus, and if noise is removed, the stimulus-driven neural correlations are highly informative about the identity of the sound or category, depending also on the duration of listening.

In the next study, we can shorten the length of the window (62.5ms) and see how successful its classification is in order to check whether the integration time of the hearing can be shorter. In addition, this time we can study other properties of the neural data than the correlation calculations. In addition to transfer learning, we can apply another network or deep learning technique for real-time design and create a human-machine interface.

## IV. CONCLUSION

We have proposed feature extraction and transfer learning methods using pre-trained neural networks for decoding aMUA signals. We have presented a comparative analysis of different pre-processed neural networks and classifiers combinations. Our results demonstrate excellently accurate decoding performance over time. The findings suggest that the correlated responses of the auditory system may play an key role in the recognition of stationary natural sounds with a shortened acquisition time.

## DECLARATION OF INTEREST

None

## Notes

http://dx.doi.org/10.6080/K03X84V3

### Competing Interest Statement

The authors have declared no competing interest.

